# Skp1 is a conserved structural component of the meiotic synaptonemal complex

**DOI:** 10.1101/2024.06.24.600447

**Authors:** Lisa E. Kursel, Kaan Goktepe, Ofer Rog

## Abstract

**Summary:** During sexual reproduction, the parental chromosomes align along their length and exchange genetic information. These processes depend on a chromosomal interface called the synaptonemal complex. The structure of the synaptonemal complex is conserved across eukaryotes, but, surprisingly, the components that make it up are dramatically different in different organisms. Here we find that a protein well known for its role in regulating protein degradation has been moonlighting as a structural component of the synaptonemal complex in the nematode *Pristionchus pacificus*, and that it has likely carried out both of these functions for more than 100 million years.

The synaptonemal complex (SC) is a meiotic interface that assembles between parental chromosomes and is essential for the formation of gametes. While the dimensions and ultrastructure of the SC are conserved across eukaryotes, its protein components are highly divergent. Recently, an unexpected component of the SC has been described in the nematode *C. elegans*: the Skp1-related proteins SKR-1/2, which are components of the Skp1, Cullin, F-box (SCF) ubiquitin ligase. Here, we find that the role of SKR-1 in the SC is conserved in nematodes. The *P. pacificus* Skp1 ortholog, Ppa-SKR-1, colocalizes with other SC proteins throughout meiotic prophase, where it occupies the middle of the SC. Like in *C. elegans,* the dimerization interface of Ppa-SKR-1 is required for its SC function. A dimerization mutant, *Ppa-skr-1^F105E^,* fails to assemble SC and is almost completely sterile. Interestingly, the evolutionary trajectory of SKR-1 contrasts with other SC proteins. Unlike most SC proteins, SKR-1 is highly conserved in nematodes. Our results suggest that the structural role of SKR-1 in the SC has been conserved since the common ancestor of *C. elegans* and *P. pacificus,* and that rapidly evolving SC proteins have maintained the ability to interact with SKR-1 for at least 100 million years.

## Introduction

The synaptonemal complex (SC) is a conserved interface that facilitates chromosome organization during meiosis. The SC aligns parental chromosomes end-to-end and regulates genetic exchanges between them, ultimately allowing for the proper segregation of chromosomes during the meiotic divisions. First identified by electron microscopy over 60 years ago, the SC is made up of two parallel axes (also called lateral or axial elements) separated by repeating striations that make up the central region of the SC (throughout, we refer to the central region of the SC simply as ‘the SC’ (Page and Hawley 2004; Zickler and Kleckner 2015)).

Despite its essential role in reproduction and its conserved ultrastructure across sexually reproducing organisms, SC components have diverged beyond recognition in multiple eukaryotic clades (Kursel, Cope, and Rog 2021; Hemmer and Blumenstiel 2016). Indeed, new SC components are still being identified, and we likely still lack the full complement of SC components in most model organisms. Further complicating molecular studies, SC components exhibit near-complete co-dependence for assembly onto chromosomes, in worms and in other organisms (Colaiácovo et al. 2003; MacQueen et al. 2002; Smolikov et al. 2007; Smolikov, Schild-Prüfert, and Colaiácovo 2009; Collins et al. 2014; Page et al. 2008; Schramm et al. 2011). Recently, co-expression of SC components allowed their purification from bacteria (Blundon et al. 2024). This suggests that SC subunits intimately associate with one another to form the repeating building blocks of an assembled SC. However, only a few intra-SC interaction interfaces have been defined (Dunce et al. 2018; Dunne and Davies 2019; Sánchez-Sáez et al. 2020; Dunce, Salmon, and Davies 2021; Kursel, Martinez, and Rog 2023), and, due to sequence divergence, it is unclear whether any of them constitute a conserved feature of the SC.

Recently, two unexpected SC proteins were identified in *C. elegans*: the Skp1-related proteins SKR-1 and SKR-2 (due to their functional redundancy we refer to them throughout as SKR-1/2; (Blundon et al. 2024)). SKR-1/2 are essential members of the Skp1, Cullin, F-box (SCF) ubiquitin ligase complex, which plays a part in virtually all eukaryotic cellular processes including germline designation (DeRenzo, Reese, and Seydoux 2003), sex determination (Clifford et al. 2000), transcriptional regulation (Ouni, Flick, and Kaiser 2010), circadian oscillation (Han et al. 2004) and hormone signaling in plants (Gray et al. 1999), to name a few (Willems, Schwab, and Tyers 2004). Within the SCF complex, Skp1 acts as an adapter by binding the N-terminus of Cul1 and the F-box motif in the F-box protein, linking the core scaffold to the substrate of the ubiquitin ligase machinery. SKR-1/2 co-purify with all other *C. elegans* SC proteins, localize to the SC, and are required for SC assembly *in vivo*. Notably, the SCF Cullin subunit CUL-1 does not localize to the SC and is not required for SC assembly. These data support the conclusion that SKR-1/2 are *bona fide* SC proteins in *C. elegans* (Blundon et al. 2024).

Here we address two outstanding questions regarding the role of SKR-1 in the SC. 1) Is the structural role of SKR-1 in the SC conserved in other nematodes? And 2) Does SKR-1 share a similar evolutionary signature to other SC proteins? We identify a single SKR-1 ortholog in the distantly related nematode *Pristionchus pacificus,* Ppa-SKR-1, and find that it localizes to the middle of the SC. Like in *C. elegans,* the predicted dimerization interface in Ppa-SKR-1 is necessary for SC assembly. Our results indicate that Ppa-SKR-1 is a structural component of the SC in *P. pacificus,* suggesting that its role in the SC originated at least 100 million years ago, in the common ancestor of *Pristionchus* and *Caenorhabditis* nematodes. Interestingly, we find that the primary sequence of SKR-1 is conserved, setting it apart from other SC proteins and shedding light on the evolutionary pressures that shape the SC.

## Results

### Identifying P. pacificus SKR-1

*C. elegans* and *P. pacificus* are a useful species pair for comparative studies. Like *C. elegans, P. pacificus* is a free-living, hermaphroditic nematode that has six pairs of chromosomes. Previous studies of meiosis in *P. pacificus* identified two SC proteins; Ppa-SYP-1 (Kursel, Cope, and Rog 2021) and Ppa-SYP-4 (Rillo-Bohn et al. 2021). Consistent with the rapid divergence of SC proteins, Ppa-SYP-4 and Ppa-SYP-1 exhibit little to no sequence homology, respectively, with their *C. elegans* counterparts. Given the recent identification of SKR-1/2 as a structural component of the SC in *C. elegans* (Blundon et al. 2024), we wondered whether SKR-1 plays a similar SC role in *P. pacificus*.

We used *C. elegans* SKR-1 as a BLASTp query against *P. pacificus* El Paco V3 predicted proteins. We identified a single strong hit which we refer to as Ppa-SKR-1. Ppa-SKR-1 clusters with *C. elegans* SKR-1/2 on a strongly supported branch to the exclusion of all other Skp1-related proteins in *P. pacificus* (Figure S1). While the *C. elegans* genome contains a recent duplication of SKR-1 called SKR-2 (Blundon et al. 2024), our phylogenetic analysis reveals that *P. pacificus* contains only one copy of SKR-1. We similarly queried seven additional *Pristionchus* proteomes and found that most species have a single SKR-1 ortholog (Figure S2)). We note that *P. pacificus*, like *C. elegans,* encodes many predicted Skp1-related proteins: 32 in *P. pacificus* and 21 in *C. elegans* (Figure S1; (Nayak et al. 2002)). While the expansion of the Skp1 family in nematodes complicates comprehensive tracing of their evolutionary history, SKR-1 orthologs appear to be the most conserved among Skp1-related proteins, and cluster together in a well-supported clade (Figure S2).

### Ppa-SKR-1 localizes to the center of the SC

We used CRISPR/Cas9 to insert an OLLAS tag on the N-terminus of Ppa-SKR-1 and examined its localization during meiosis (Figure 1). OLLAS::Ppa-SKR-1 appears as threads on meiotic chromosomes from the time of SC assembly at meiotic entry, throughout pachytene (the stage when the SC is completely assembled on all chromosomes), and to diplotene (the extended stage of SC disassembly; Figure 1A). This pattern matches that of other SC proteins (Rillo-Bohn et al. 2021; Kursel, Cope, and Rog 2021). The axis component HOP-1 (Rillo-Bohn et al. 2021) localizes to meiotic chromosomes slightly before OLLAS::Ppa-SKR-1 as faint lines indicative of unpaired chromosomes (Figure 1B). As OLLAS::Ppa-SKR-1 signal begins to overlap with HOP-1, the lines of HOP-1 are brighter, reflecting paired, synapsed chromosomes. During diplotene, OLLAS::Ppa-SKR-1 remains on the bright-staining regions of HOP-1 until the SC fully disassembles.

**Figure 1:**
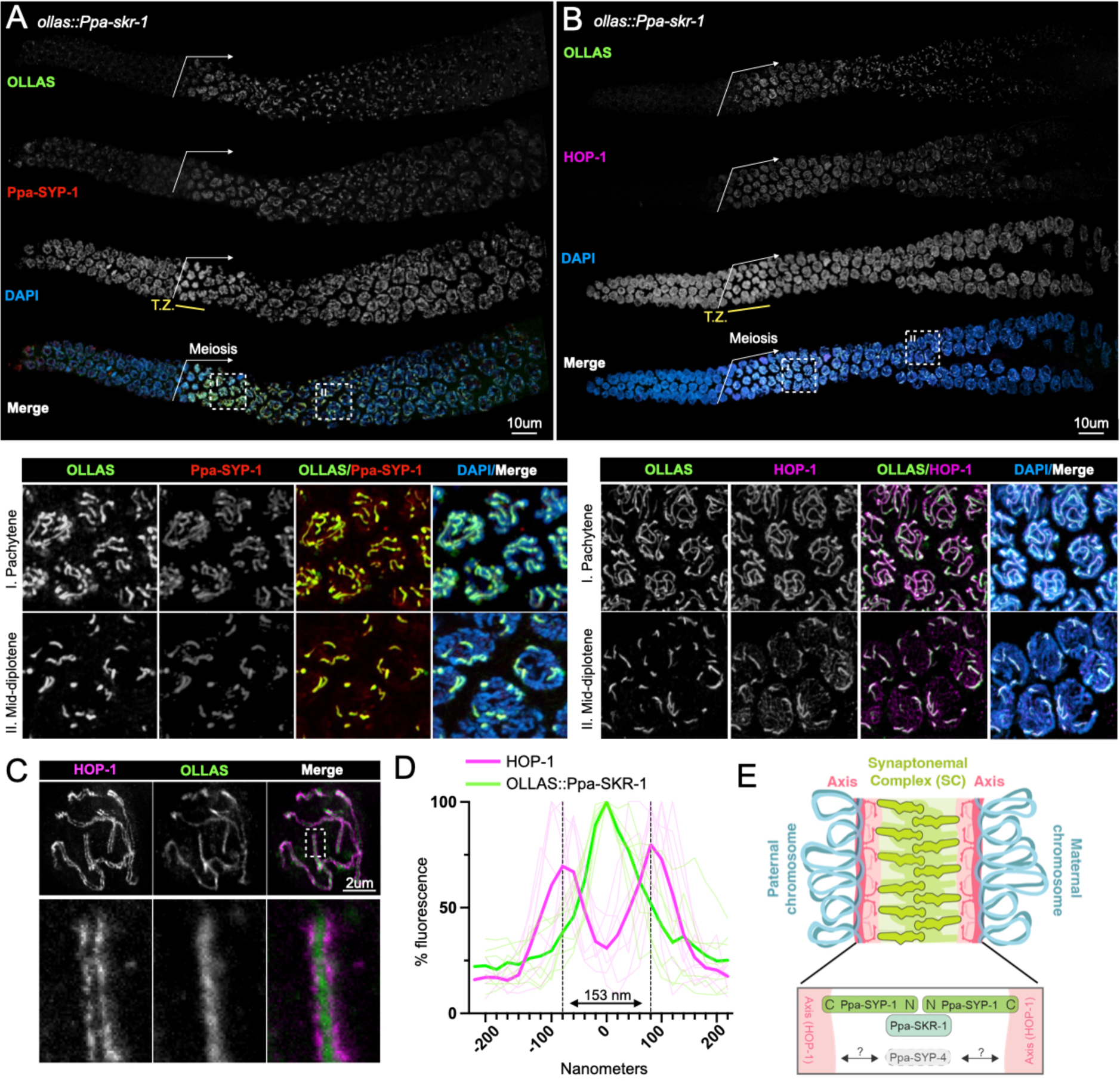
Ppa-SKR-1 localizes to the middle of the SC. (A) Top panel, confocal image of whole gonads from *ollas::Ppa-skr-1* stained with anti-OLLAS, anti-SYP-1 and DAPI. Bottom panel, zoom in on pachytene (I) or mid-diplotene (II) nuclei. (B) Confocal image as in (A) except with HOP-1 staining. In (A) and (B), the beginning of the meiotic gonad is indicated with a white arrow and the transition zone is labeled below the DAPI channel in yellow (T.Z.). (C) Super-resolution STED image of a single pachytene nucleus from *ollas::Ppa-skr-1* worms stained with anti-OLLAS and anti-HOP-1. Zoom-in panels show OLLAS::Ppa-SKR-1 between parallel HOP-1 tracks. (D) Plot of line scans of pixel intensity for anti-HOP-1 and anti-OLLAS across parallel axes in *ollas::Ppa-skr-1* worms. The average distance between parallel axes is 153nm. (E) Cartoon of the *P. pacificus* synaptonemal complex with the orientation and position of Ppa-SYP-1 and Ppa-SKR-1 relative to HOP-1 indicated in the bottom panel. The relative position of Ppa-SYP-4 is not known (grey arrows and question marks). Adapted from (Kursel, Aguayo Martinez, and Rog 2023).

SC proteins occupy stereotypical positions in the ∼150nm space separating the two parental chromosomes. Ppa-SYP-1, like its *C. elegans* counterpart, spans the 100nm width of the SC in a head-to-head manner (N-terminus in, C-terminus out) such that staining with a C-terminal epitope produces two parallel lines and N-terminal staining produces a single thread in the middle of the SC (Köhler et al. 2020; Kursel, Cope, and Rog 2021; Schild-Prüfert et al. 2011). Using STED super-resolution microscopy, we found that the axis protein HOP-1 formed parallel tracks that are 153nm wide on average (Figure 1C, D) and that Ppa-SKR-1 localized to the central region of the SC, midway between the parallel HOP-1 tracks. These cytological data indicate that, like in *C. elegans*, Ppa-SKR-1 occupies the middle of the SC ladder, where the N-terminus of SYP-1 is located (Figure 1E, (Blundon et al. 2024)).

### The Ppa-SKR-1 dimerization interface is required for SC assembly

The essential functions of Skp1 make it challenging to study its role in the SC. *C. elegans* worms lacking both SKR-1 and −2 fail to hatch, reflecting the essential roles of SCF in embryogenesis and cell proliferation (Nayak et al. 2002; Blundon et al. 2024). Given that *P. pacificus* harbors a single Skp1 ortholog, we predicted that gene deletion would result in embryonic lethality. We therefore wished to generate a separation-of-function allele of *Ppa-skr-1*.

Previous studies found that Skp1 dimerizes *via* a conserved hydrophobic interface that is not essential for F-box binding (Kim et al. 2020; Henzl, Thalmann, and Thalmann 1998). In *C. elegans,* mutations that disrupt SKR-1/2’s ability to dimerize (*skr-1^F115E^*) cause a complete failure of SC assembly and prevent SKR-1/2 localization to an already formed SC. Importantly, these mutations do not abolish SCF activity, suggesting that SKR-1/2 dimerization is necessary specifically for SC function (Blundon et al. 2024).

We used structural homology to predict the dimerization interface in Ppa-SKR-1 (Figure 2A). We found that a residue critical for dimerization in *Dictyostelium* Skp1, F97 (Kim et al. 2020), aligns closely with F105 in Ppa-SKR-1 (Figure 2A). To test the function of the putative dimerization interface, we used CRISPR/Cas9 to make *ollas::Ppa-skr-1^F105E^.* Gratifyingly, we easily obtained *ollas::Ppa-skr-1^F105E^* homozygous animals. Out of 46 F2s singled from heterozygous *ollas::Ppa-skr-1^F105E^*F1 parents, 12 were homozygous wildtype, 22 were heterozygous and 12 were homozygous for *ollas::Ppa-skr-1^F105E^,* matching expected Mendelian ratios. This suggests that the F105E mutation does not disrupt SCF functions during development.

**Figure 2:**
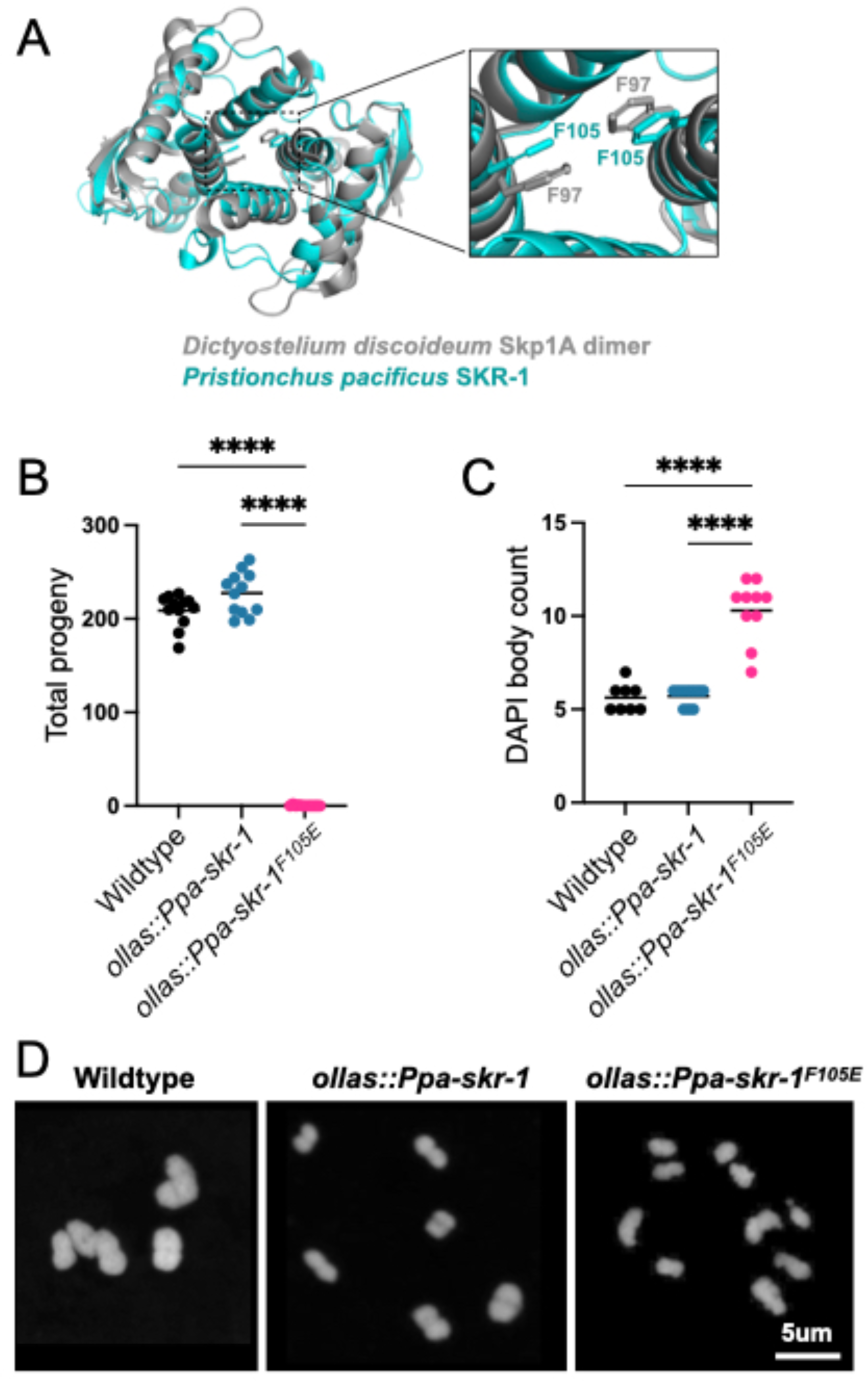
Conserved dimerization interface in SKR-1 is required for *P. pacificus* meiosis. (A) Alignment of *P. pacificus* SKR-1 AlphaFold model (cyan) to *Dictyostelium* Skp1A dimer NMR structure (PDB structure 6V88, gray). Conserved phenylalanines required for dimerization are labeled in zoom. Dot plot depicting total progeny (B) and DAPI body count (C) for wild-type *P. pacificus, ollas::Ppa-skr-1 and ollas::Ppa-skr-1^F105E^.* Asterisks reflect P-values from Tukey’s multiple comparison test where **** indicates P < 0.0001. (D) Representative images of DAPI-stained Meiosis I bivalents (DAPI bodies) from the indicated genotypes.

To evaluate successful completion of meiosis, we counted total progeny in wild-type, *ollas::Ppa-skr-1* and *ollas::Ppa-skr-1^F105E^*worms. Total progeny produced by *ollas::Ppa-skr-1* worms were comparable to that of the wild-type *P. pacificus*, indicating that the OLLAS insertion did not interfere with meiosis. In contrast, *ollas::Ppa-skr-1^F105E^*worms were almost sterile, mimicking other SC null mutants (Figure 2B). Notably, several homozygous hermaphrodites produced one to two progeny, further indicating that OLLAS::Ppa-SKR-1^F105E^ can carry out the non-meiotic functions of Skp1 proteins. Together, this analysis indicated that Ppa-SKR-1 dimerization is necessary for reproduction.

To examine meiotic dysfunction in more detail, we monitored successful formation of crossovers in meiotic prophase. Chromosomes that form a crossover are joined at metaphase of Meiosis I, forming so-called “bivalents” that can be visualized by staining DNA with DAPI. Since *P. pacificus* has six chromosome pairs, successful generation of a crossover on each pair yields six DAPI-staining bodies. We found no significant difference in DAPI body counts between wild-type and *ollas::Ppa-skr-1* worms. They averaged 5.6 and 5.7 DAPI bodies, respectively (Figure 2C). However, *ollas::Ppa-skr-1^F105E^* worms had a significantly elevated DAPI body count (mean = 10.5) suggesting that failure of chromosome pairing or crossover formation underlies the reduced progeny count in ollas::*Ppa-skr-1^F105E^*worms (Figure 2D).

Cytological examination established that *Ppa-skr-1^F105E^* worms lack an SC. Meiotic nuclei in the mutant spent an extended duration in the transition zone - the region of the gonad where the SC assembles, marked by crescent-shaped nuclei (Figure 3, compare to Figure 1A, B). An increase in transition zone length is seen in other SC mutants (MacQueen et al. 2002; Colaiácovo et al. 2003; Smolikov et al. 2007; Smolikov, Schild-Prüfert, and Colaiácovo 2009) and is thought to reflect a synapsis checkpoint (Harper et al. 2011). HOP-1 appeared as thin tracks throughout the gonad in *Ppa-skr-1^F105E^* worms, indicative of chromosomes that were unable to assemble an SC (Figure 3B). Furthermore, Ppa-SYP-1 staining revealed complete lack of SC assembly (Figure 3C). In *C. elegans* and other species, SC components seem to be required for each other’s stability (Colaiácovo et al. 2003; Hurlock et al. 2020; Smolikov et al. 2007; Smolikov, Schild-Prüfert, and Colaiácovo 2009; Blundon et al. 2024; Z. Zhang et al. 2020). Indeed, Ppa-SYP-1 staining was almost completely absent in *ollas::Ppa-skr-1^F105E^*worms. Moreover, when SC components are present but cannot load onto chromosomes, SC material forms large aggregates called polycomplexes (Page and Hawley 2004). Notably, polycomplexes are absent in *ollas::Ppa-skr-1^F105E^* worms (Figure 3) and in *C. elegans skr-1^F115E^*worms (Blundon et al. 2024), suggesting other SC component are not able to assemble in the dimerization mutant. These data indicate that, like in *C. elegans,* SC formation in *P. pacificus* depends on Ppa-SKR-1 dimerization. Taken together with Ppa-SKR-1 localization (Figure 1), our data indicate that Ppa-SKR-1 is a structural component of the *P. pacificus* SC.

**Figure 3:**
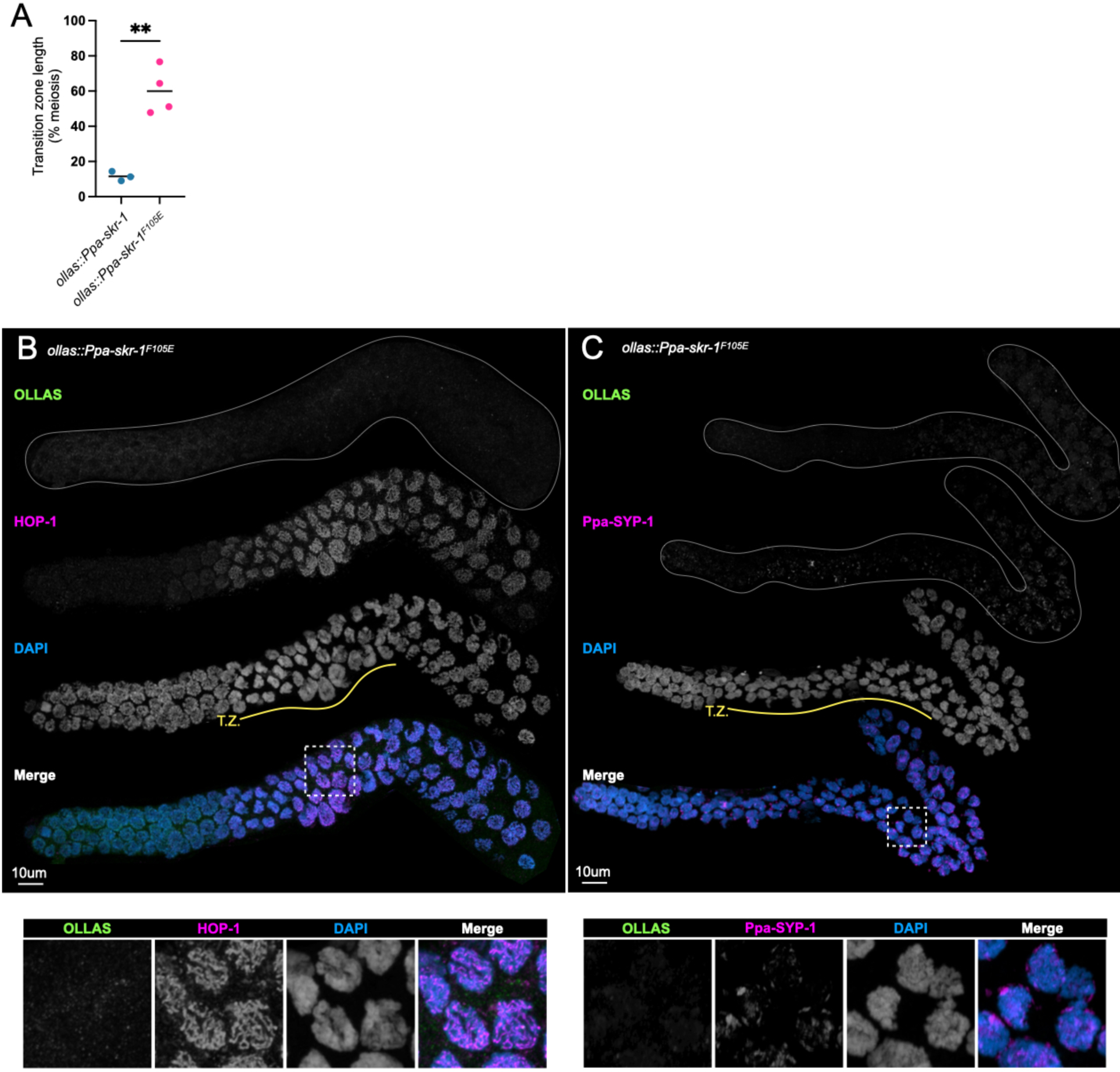
Ppa-SKR-1^F105E^ fails to assemble the SC. (A) Dot plot showing transition zone length as percent of meiosis. Asterisks reflect the P-value from an unpaired T-test where ** indicates P < 0.01. (B) and (C), Confocal images of whole gonads from *P. pacificus ollas::skr-1^F105E^* stained with anti-OLLAS, anti-HOP-1 (B) or anti-SYP-1 (C), and DAPI. Lower panels in (B) and (C) show zoom-in on regions indicated by white, dashed boxes and the transition zone is labeled below the DAPI channel in yellow (T.Z.).

### Unlike other SC components, SKR-1 sequence is conserved in nematodes

We previously found that SC proteins in nematodes*, Drosophila* and mammals have a unique evolutionary signature; diverged protein sequence but conserved length and position of coiled-coil domains and conserved overall protein length (Kursel, Cope, and Rog 2021). We hypothesized that this evolutionary signature could be explained by the SC mode of assembly, which likely relies on weak multi-valent interactions mediated by coiled-coil domains. Since the sequence requirements for coiled-coil domains are flexible (typically defined as a heptad repeat where the first and fourth residues are hydrophobic and the fifth and seventh are charged or polar), selection to maintain coiled-coil domains could allow for significant sequence divergence. At the time of our analysis, SKR-1 had not been identified as an SC protein. Therefore, we wished to compare the evolutionary signature of SKR-1 to the other SC proteins.

Unlike the other SC proteins in *Caenorhabditis* and *Pristionchus,* the sequence of SKR-1 is conserved in both clades, ranking in the bottom one percentile for amino acid substitutions per site (Figure 4A). Unsurprisingly, residues involved in CUL-1 binding, F-box protein binding, and the dimerization interface are highly conserved, even between *C. elegans, P. pacificus* and *H. sapiens* (Figure 4B). We also found that SKR-1 does not contain conserved coiled-coil domains (Figure 4C, S3A). *Pristionchus* SKR-1 does have a low-scoring predicted coiled-coil domain from amino acids 20 – 47 (Figure S3A). However, AlphaFold does not predict a coiled-coil formed by Ppa-SKR-1 and this coiled-coil signature is not conserved in *Caenorhabditis* (Figure S3A) or in *Dictyostelium*, where the corresponding residues are mostly disordered in the NMR structure (Kim et al. 2020). Together, this argues against the functional importance of coiled-coil domains in SKR-1 (Figure S3B). Lastly, the length of SKR-1 is conserved, like other SC proteins (Figure 4D). Taken together, our analysis indicates that the evolutionary trajectory of SKR-1 is distinct from other SC proteins in *Caenorhabditis* and *Pristionchus* and suggests that its interaction with other SC proteins is mediated by domains other than coiled-coils.

**Figure 4:**
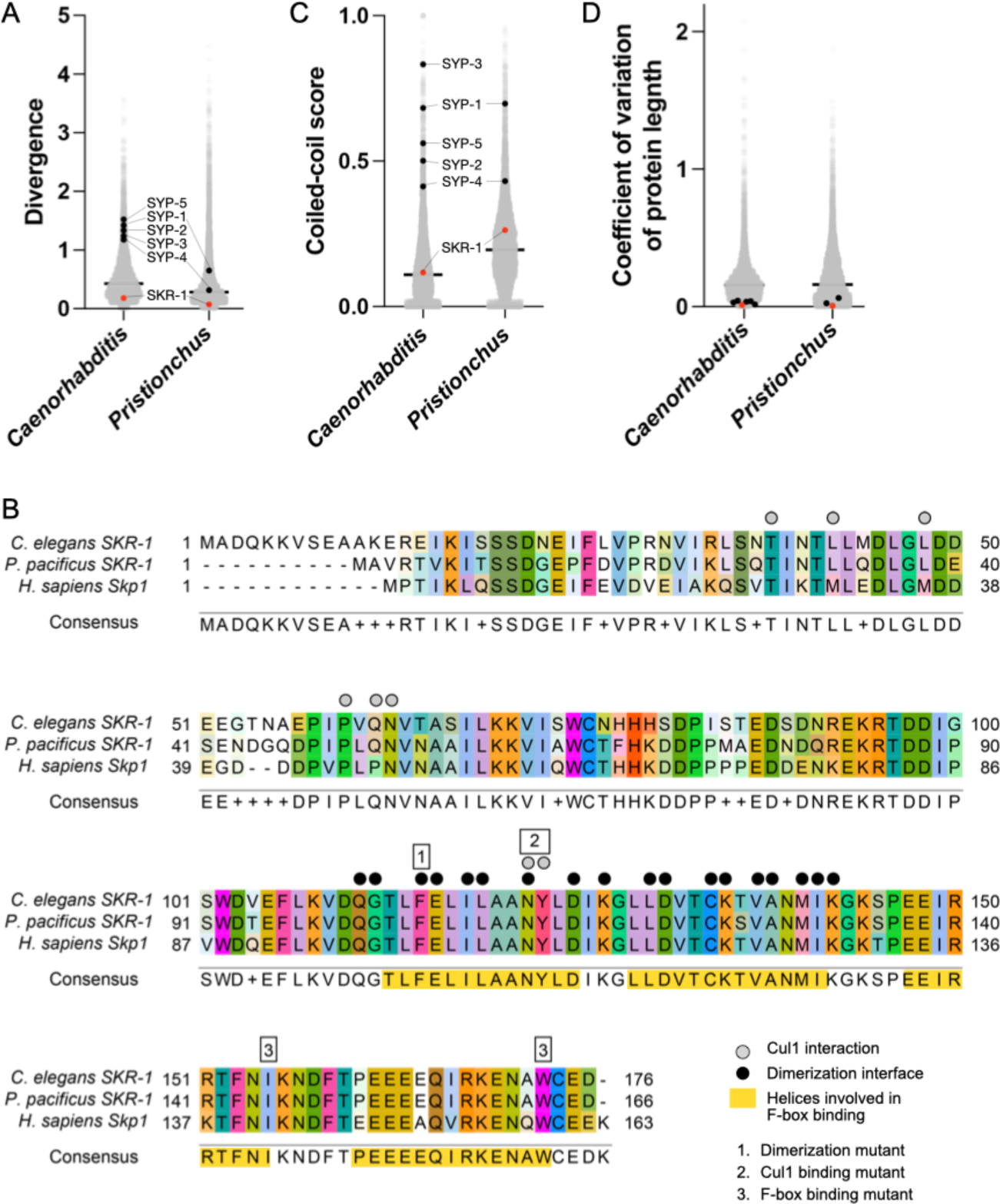
SKR-1 has an evolutionary signature distinct from other SC proteins. (A) Dot plot showing protein divergence for the *Caenorhabditis* and *Pristionchus* proteomes. SYP proteins and SKR-1 are indicated (black and pink, respectively). (B) Alignment of Skp1 orthologs from *C. elegans* and *P. pacificus,* and *H. sapiens* with Cul1 interaction, dimerization and F-box binding sites labeled (Zheng et al. 2002; Kim et al. 2020). Additionally, three mutants generated by Blundon and Caesar *et al*. are indicated by numbered boxes (Blundon et al. 2024). (C) and (D), dot plots showing coiled-coil conservaBon and coefficient of variaBon of protein length for the *Caenorhabdi+s* and *Pris+onchus* proteomes. SYP proteins and SKR-1 are indicated as in (A).

## Discussion

We found that SKR-1 is a structural member of the SC in *P. pacificus.* Ppa-SKR-1 dynamically localizes to meiotic chromosomes in a manner that is indistinguishable from that of other SC proteins. Like other SC proteins, Ppa-SKR-1 exhibits stereotypic localization relative to the axes: it localizes to the middle of the SC, placing it near the N-terminus of Ppa-SYP-1 (Figure 1E). Finally, like in *C. elegans,* the dimerization interface of Ppa-SKR-1 is necessary for SC assembly but not for other essential functions. Taken together, our cytological, functional and phylogenetic data suggest that the function of SKR-1 as a structural component of the SC has been conserved since the common ancestor of *C. elegans* and *P. pacificus*, at least 100 million years ago.

Our work on the conservation of an SC role for SKR-1 in nematodes raises the possibility that it extends to Skp1 proteins in other clades. Unsurprisingly, proteasome-mediated degradation regulates multiple key steps in meiosis (Ahuja et al. 2017; Rao et al. 2017; Guan et al. 2022) and the proteasome itself localizes to the SC in *C. elegans* and mice (Rao et al. 2017; Ahuja et al. 2017). Skp1 also localizes to the SC in male and female mice (Guan et al. 2020), and in *Arabidopsis* plants where it is called ASK1 (Wang et al. 2004). In both cases, its disruption leads to meiotic defects (Yang et al. 2006). However, the essential functions of the proteosome and Skp1, and the consequent far-ranging effects of their disruption, has made it difficult to parse their role in the protein degradation from any potential structural role in the SC.

*C. elegans* has proved to be an especially valuable system for studying the role of Skp1 in the SC because it contains two partially redundant paralogs, SKR-1 and SKR-2. Having two SKR-1 paralogs allowed Blundon and Cesar *et al*. to identify the separation-of-function dimerization mutant. We similarly found that a mutation predicted to disrupt Ppa-SKR-1 dimerization results in separation of function; worms are viable and have no obvious growth defects indicating SCF functions are intact, but they are sterile due to failure of SC assembly. It will be interesting to explore whether the corresponding Skp1 dimerization interface - which is conserved at the protein sequence level in mammals and plants - would help to generate separation-of-function alleles in other model organisms.

The molecular details of SKR-1 interaction with other SC components remain unknown in both *C. elegans* and *P. pacificus*. SKR-1 proteins are not merely recruited to the SC like other so-called ‘client’ proteins, including the crossover regulator family ZHP-1/2/3/4 (Jantsch et al. 2004) and the polo-like kinase PLK-2 (L. Zhang et al. 2018; Harper et al. 2011; Labella et al. 2011). For example, the localization pattern of ZHP-1/2/3/4 is distinct from SC proteins and the SC can still assemble in the absence of the ZHPs. In contrast, SKR-1 is essential for SC assembly in both *C. elegans* and *P. pacificus*, and it contributes to the stability of SC components *in vivo* and *in vitro*. Such intimate co-dependence suggests the existence of underlying protein-protein interactions that provide specificity and stability.

The protein surfaces that mediate interactions between SC proteins must co-evolve to maintain compatibility. In this light, the high conservation of SKR-1 *versus* the high divergence of other SC components might seem surprising since proteins in complex often have homogenous evolutionary rates (Wong et al. 2008) and genes whose evolutionary rates covary tend to be functionally related (Clark, Alani, and Aquadro 2012). However, a more recent study reported that direct physical interaction is only a weak driver of evolutionary rate covariation (Little, Chikina, and Clark 2024). Moreover, moonlighting proteins that function in multiple complexes can confound such analyses. Taking these factors into account, SKR-1’s role in the highly conserved SCF complex might overwhelm any signal of shared evolutionary rates with other SC proteins. In addition, we note that the SC is a condensate (Rog, Köhler, and Dernburg 2017), and that many condensates rely on weak, multivalent interactions to recruit and exclude member and non-member components, respectively (Shin and Brangwynne 2017). SC proteins might have multiple, redundant interaction interfaces with SKR-1, each too weak to pose a strong constraint on the primary sequence.

The recent duplication of SKR-1 in the lineage leading to *C. elegans* (Blundon et al. 2024) could suggest that gene duplication has allowed Skp1 proteins to adopt a novel function - a structural component of the SC. However, our findings suggest that the role of SKR-1 in the SC is more ancient and that a single SKR-1 protein has likely performed both functions in the common ancestor of *C. elegans* and *P. pacificus*. An ancestral dual-function protein suggests that SKR-1 has been subjected to evolutionary pressures to maintain both functions for at least 100 million years. Interestingly, SKR-1’s dual roles in SCF and the SC entail that mutations in *skr-1* might have pleiotropic effects in development (SCF) *versus* reproduction (SC). If so, *C. elegans* may represent a lineage where such intralocus conflict is resolving by gene duplication and specialization (Castellanos, Wickramasinghe, and Betrán 2024). In this scenario, the different structural and functional requirements of the SC *versus* the SCF complex could be divided between SKR-1 and SKR-2, allowing them to eventually differentiate into an SC-dedicated protein and an SCF-dedicated one. Such specialization has likely taken place throughout the broader Skp1-related gene family, which has massively expanded in nematodes (Nayak et al. 2002). Intralocus conflict and related processes provide a leading framework in the evolution of aging (Adler and Bonduriansky 2014), suggesting that the evolutionary trajectory of SKR-1 in nematodes could shed light on the evolution of aging more broadly.

## Materials and Methods

### Worm strains and maintenance

We used *Pristionchus pacificus* strain PS312 for the wildtype control and for injections to make *ollas::Ppa-skr-1.* To make *ollas::Ppa-skr-1^F105E^,* we injected into *ollas::Ppa-skr-1.* All strains were grown at 20°C on NGM agar with OP50 bacteria. We maintained PS312 and *ollas::Ppa-skr-1* in a homozygous state but since *ollas::Ppa-skr-1^F105E^* was sterile, we maintained it as a heterozygous line by singling animals and genotyping each generation. We consistently observed severe SC defects in one-quarter of the progeny from a heterozygous parent and never observed severe defects in progeny from *ollas::Ppa-skr-1* or PS312 parents. For DAPI body counts, we identified gonads with SC defects in progeny of *ollas::Ppa-skr-1^F105E^* heterozygous animals, and considered those gonads with severe SC defects to be homozygous. To perform progeny counts of *ollas::Ppa-skr-1^F105E^*, we singled from a heterozygous parent, counted progeny and genotyped by PCR (see below) after the complete brood was laid.

### Identification of P. pacificus SKR-1

We used *C. elegans* SKR-1 as a query in a BLASTp search, implemented on pristionchus.org, of the *P. pacificus* El Paco V3 genome (Dieterich et al. 2007). The top hit was ppa_stranded_DN29817_c0_g1_i2, a 166 amino acid protein. We also performed a tBLASTn search using *C. elegans* SKR-1 as a query against the El Paco V4 genome (GCA_000180635.4) implemented on ncbi.nlm.nih.gov. This identified the coding sequence KAF8362560.1, which encodes a 166 amino acid protein identical to ppa_stranded_DN29817_c0_g1_i2. When we used the 166 amino acid protein as a query in a BLASTp search of the *C. elegans* proteome, the top his was *C. elegans* SKR-1 (F46A9.5).

We note that performing the same BLASTp search against the *P. pacificus* genome on wormbase.org (Sternberg et al. 2024) produces a top hit to PPA23980, a protein with 1443 amino acids that contains a predicted ABC transporter transmembrane domain in its N-terminus and homology to SKR-1 in its C-terminus. We suspect that this is due to an annotation error that merges two genes since wormbase.org also hosts the El Paco V4 genome assembly and the start codon of the 166 amino acid version of SKR-1 is preserved in PPA23980.

To confirm that ppa_stranded_DN29817_c0_g1_i2 is indeed the SKR-1 ortholog in *P. pacificus*, we generated a neighbor-joining phylogenetic tree with all hits that resulted from BLASTp search of *P. pacificus* with *C. elegans* SKR-1 (File S1, S2, S3). Since *P. pacificus* ppa_stranded_DN29817_c0_g1_i2 groups closest with *C. elegans* SKR-1/2 (Figure S1, File S3), it is most likely to be the SKR-1 ortholog. Thus, we refer to ppa_stranded_DN29817_c0_g1_i2 as Ppa-SKR-1.

### Sequence collection, alignment and phylogenetic analysis

We identified *Caenorhabditis* SKR-1 orthologs using the EnsEMBL Compara pipeline implemented on wormbase.org (Harris et al. 2010). We only kept sequences from species with a single predicted ortholog, with the exception of *C. elegans,* which has an SKR-1 paralog, SKR-2, leaving 16 SKR-1 sequences for analysis. We identified *Pristionchus* SKR-1 orthologs by performing BLASTp with *C. elegans* SKR-1 against the eight *Pristionchus* genomes available on Pristionchus.org (Dieterich et al. 2007). We saved the top hit from each search. We used Clustal Omega for all protein alignments and Geneious Tree Builder (neighbor-joining method, Geneious Prime version 2023.2.1) with 100x bootstrap resampling to generate the phylogenies in supplementary Figures 1 and 2. All protein sequences, alignments and trees are available as supplemental data (File S4 – S9).

### CRISPR genome editing

We aimed to insert an OLLAS tag in the N-terminus of Ppa-SKR-1, immediately following the start methionine. We made an injection mix containing 1ul Cas9 (IDR, Alt-R S.p. Cas9 Nuclease V3, 10ug/ul), 3.5ul repair template (200uM), 3.5ul annealed tracr/crRNA mix and 0.5ul duplex buffer (IDT). We injected the gonads of wildtype (PS312) young adult hermaphrodite *P. pacificus* and singled each injected worm to its own plate. We extracted DNA from ∼16 combined F1 worms from each plate and genotyped with primers that span the OLLAS insertion site (LEK1094 GTTTCACAACAACGGCCCTC and LEK1095 CTTGATGACGTCACGGGGAA) to identify “jackpot plates” (*i.e.,* plates with high rates of OLLAS insertion). We singled as many F1s as possible from the jackpot plates and genotyped again to identify individual insertion events.

To make *ollas::Ppa-skr-1^F105E^* we followed a similar strategy as above except we injected into *ollas::Ppa-skr-1.* We screened the pooled F1s by doing PCR with primers LEK1111 (GAGAAGGGAACAACGTGGGT) and LEK1112 (CGCGCGTCTCATTCAACAAA) and digesting with MboI. The predicted Cas9 cut site is near an MboI site in *ollas::skr-1,* so CRISPR repair events could destroy the MboI site. In this scenario, wildtype plates will have bands that are 259, 241 and 92 base pairs in length after MboI digest but plates that contain CRISPR mutants will also have a 351 base pair band. We singled F1s from plates with the 351 base pair band and did a second round of genotyping with LEK1111 and LEK1112, this time followed by digest with SalI. Animals that contain CRISPR repair events from the injected homology template will gain an SalI site. PCR from wildtype animals will remain undigested (592 base pairs) whereas PCR from a mutant animal will get cut (336 and 256 base pair bands). See Table S1 for a list of primers, crRNAs and repair templates used for CRISPR.

### Immunofluorescence and confocal microscopy

We prepared gonads for immunofluorescence and confocal microscopy as we have done previously (Kursel, Cope, and Rog 2021; Phillips, McDonald, and Dernburg 2009). Briefly, we dissected age-matched adult worms in egg buffer with 0.01% Tween-20 and fixed in a final concentration of 1% formaldehyde. We transferred samples to a HistoBond microscope slide, froze for 1 minute on dry ice and quickly immersed the slide in −20°C methanol for one minute. Slides were washed in PBST and blocked in Roche Block (Cat # 11096176001) for 30 minutes at room temperature. We incubated the slides in 80 µl of primary antibody overnight at 4°C. Primary antibody concentrations were as follows: Rabbit anti-PPA-SYP-1 1:500 (Kursel, Cope, and Rog 2021), Rat anti-OLLAS 1:200 (Invitrogen Catalog # MA5-16125), Rabbit anti-PPA-HOP-1 1:300 (Rillo-Bohn et al. 2021). The following day, slides were washed for three rounds of 10 minutes in PBST, then incubated in secondary antibody. Secondary antibody concentrations were as follows: Donkey anti-rabbit Cy3 1:500 and Donkey anti-rat Alexa488 1:500 (Jackson ImmunoResearch). Finally, we washed slides in PBST and DAPI and mounted them in NPG-Glycerol. Slides were imaged on a Zeiss LSM880 confocal microscope with Airyscan and a 63 × 1.4 NA oil objective. Confocal images presented in this manuscript are maximum intensity projections.

### STED super-resolution microscopy

Gonads for STED microscopy were prepared as for confocal microscopy with the following changes: 1) we omitted DAPI staining, 2) we used Goat anti-Rabbit STAR RED 1:200 (Abberior # STRED-1002-500UG) and Goat anti-Rat Alexa 594 1:200 (Jackson ImmunoResearch) as secondaries, and 3) we mounted the samples in Abberior Mount Liquid Antifade (Abberior # MM-2009-2X15ML). Samples were imaged on Aberrior STEDYCON mounted on a Nikon Eclipse Ti microscope with a 100 × 1.45 NA oil objective. Line scans were performed in FIJI (Schindelin et al. 2012).

### Structural modeling and alignment

We used AlphaFold (Jumper et al. 2021), implemented in ColabFold (Mirdita et al. 2022), to model the structure of full-length Ppa-SKR-1. We used Pymol ((Schrodinger 2015), version 2.5.7) to visualize Ppa-SKR-1 and to align it to the *Dictyostelium* Skp1A dimer NMR structure ((Kim et al. 2020), PDB structure 6V88).

### Progeny counts

We singled twelve L4s from each genotype and grew them at 20°C. We moved the parents to a fresh plate every day for four days and counted the progeny after allowing them to mature for up to five days. For the *ollas::Ppa-skr-1^F105E^* genotype, we singled 50 F1s from a heterozygous animal. We moved the F1s to fresh plates daily as described. At the end of the fourth day of egg laying, we identified the homozygous animals among the F1s by genotyping the parent with LEK1111/LEK1112 PCR primers followed by SalI digest as described above. We counted progeny from those animals confirmed to be homozygous mutants.

### Calculating divergence, coiled-coil conservation and length conservation

The *Caenorhabditis* and *Pristionchus* proteome values (Figure 4A, 4C and 4D) were published previously (Kursel, Cope, and Rog 2021). We calculated divergence values, coiled-coil conservation scores and coefficient of variation of protein length for SKR-1 as we have done previously for SC proteins (Kursel, Cope, and Rog 2021) using SKR-1 orthologs from *Caenorhabditis* or *Pristionchus* collected as described above.

### Statistical analysis

We used an ordinary one-way ANOVA with Tukey’s multiple comparisons test to test for differences in total progeny and DAPI body counts between genotypes (Figure 2B and 2C). In Figure 3A, we used an unpaired t test to test for differences in transition zone length.

## Data availability

Worm strains generated in this study are available by request. All sequence alignments and phylogenies are included as supplementary data files. Proteome-wide analysis of divergence, coiled-coil scores and protein length variation in *Caenorhabditis* and *Pristionchus* was published previously (Kursel, Cope, and Rog 2021).

## Acknowledgements

We would like to thank the Rog Lab for discussions, Abby Dernburg for antibodies and Yumi Kim for discussions and for sharing data prior to publication. Some strains used in this work were provided by the CGC, which is funded by NIH Office of Research Infrastructure Programs (P40 OD010440). We acknowledge the HSC Cell Imaging Core at the University of Utah for use of the STED microscope and The University of Utah Center for High Performance Computing for computational resources. KG was supported by the Undergraduate Research Opportunity Program at the University of Utah and by the Biology Research Scholar Award from the School of Biological Sciences. This work was supported by grants R35GM128804 from NIGMS and 2219605 from NSF.

---

**Figure S1:**
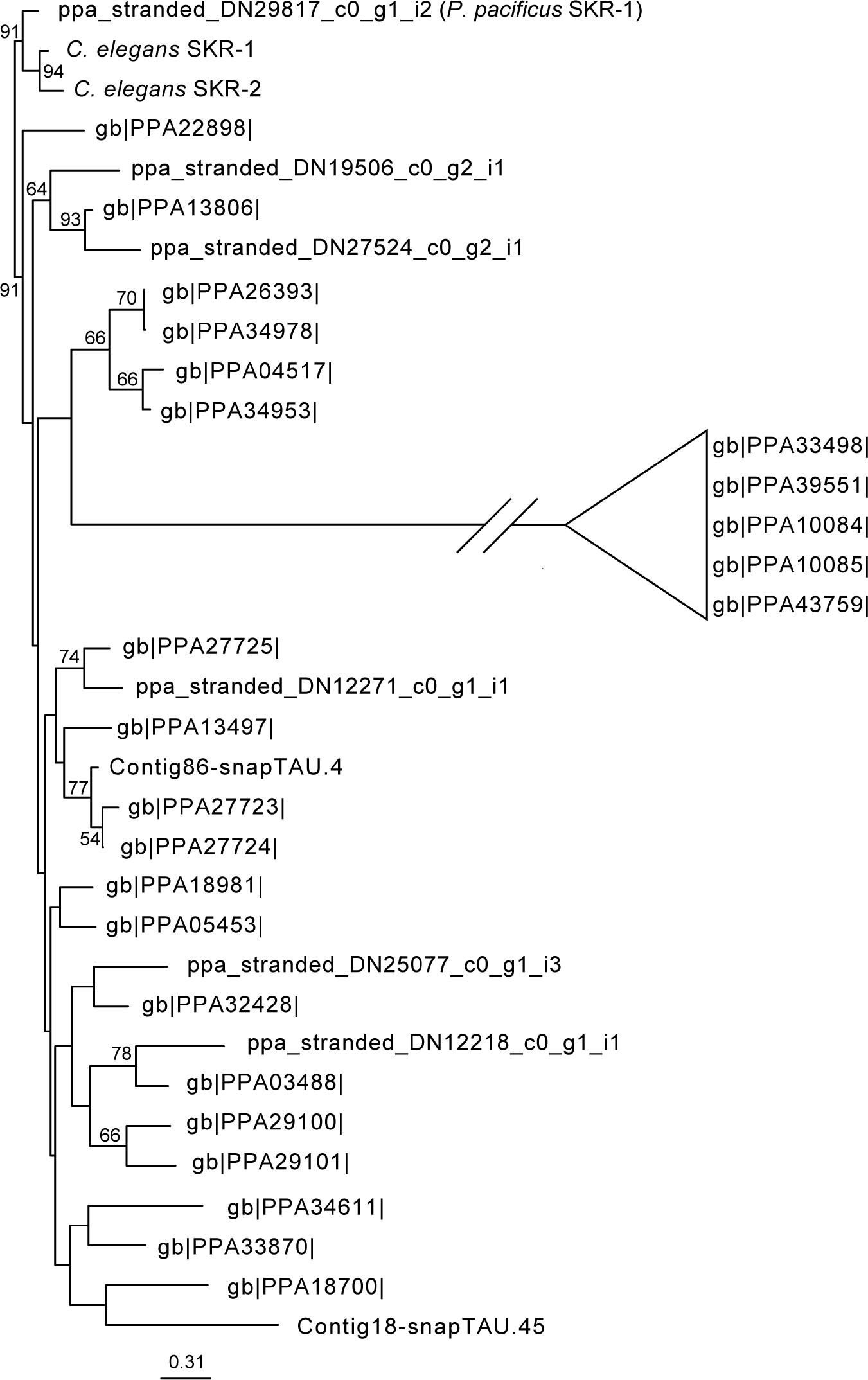
Neighbor-joining phylogenetic tree of *P. pacificus* Skp1-related proteins. Phylogenetic tree made from a protein alignment of all *P. pacificus* Skp1-related proteins identified *via* BLASTp search. Bootstrap values greater than 50 are displayed. Note: the branch leading to PPA33498, PPA39551, PPA10084, PPA10085 and PPA43759 was truncated (diagonal lines) to more easily display the entire phylogeny.

**Figure S2:**
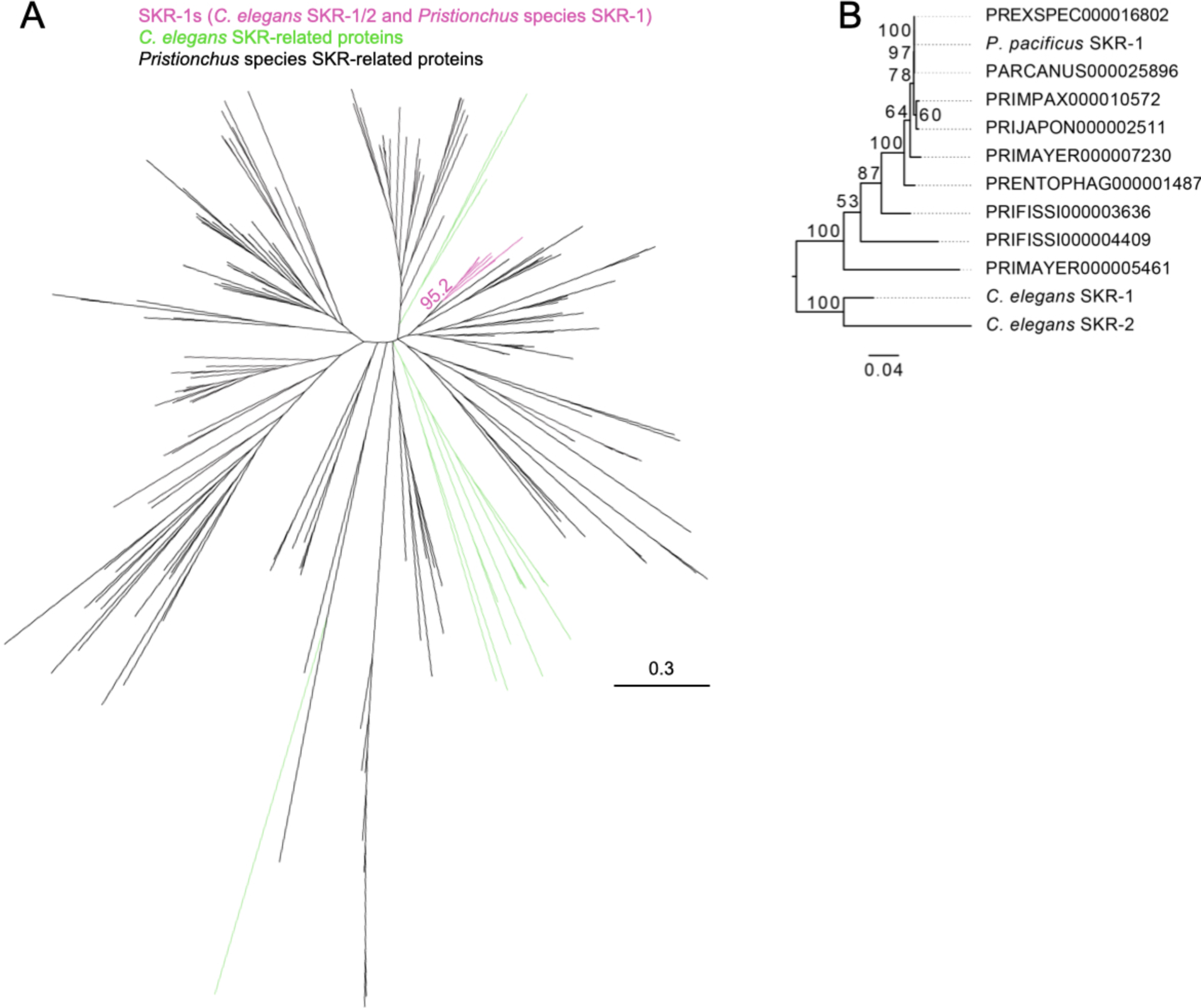
Neighbor-joining phylogenetic tree of *Pristionchus* Skp1-related proteins. (A) Unrooted phylogenetic tree with 100x bootstrap support made from a protein alignment of all Skp1-related proteins from *C. elegans* and eight *Pristionchus* species. The clade containing *C. elegans* SKR-1/2 and *P. pacificus* SKR-1 has pink branches, all other *C. elegans* SKRs have green branches and all other *Pristionchus* Skp1-related proteins have black branches. The bootstrap support value for the SKR-1 clade is shown. (B) Phylogenetic tree with 100x bootstrap support made from an alignment of the proteins in the SKR-1 clade in (A, pink branches). The tree is rooted on the common ancestor of *Caenorhabditis* and *Pristionchus*.

**Figure S3:**
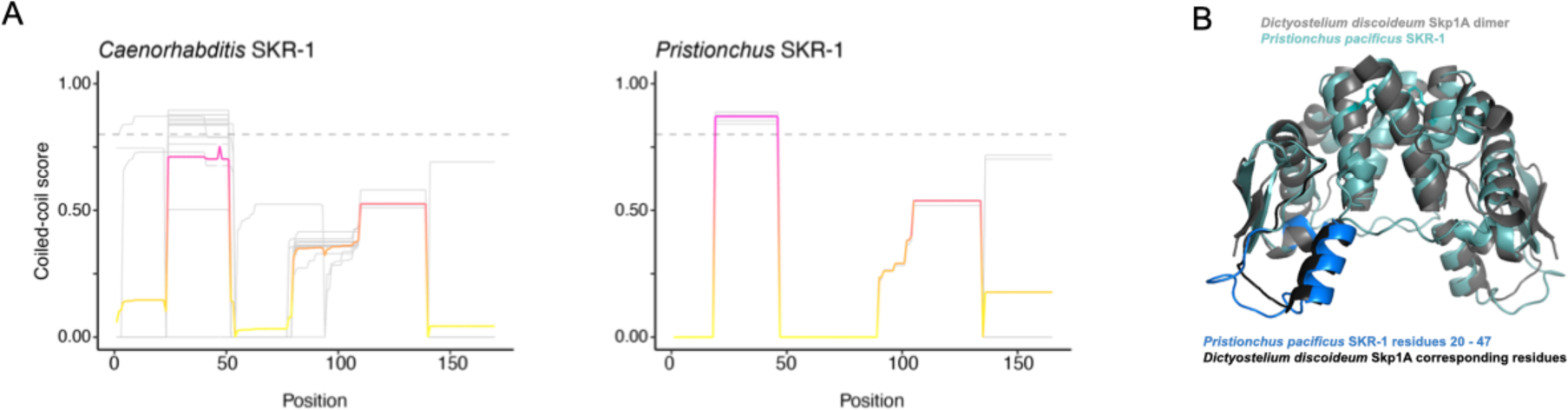
SKR-1 does not contain conserved coiled-coil domains. (A) Plot showing likelihood of coiled-coil domain at every residue in *Caenorhabditis* and *Pristionchus* SKR-1. Individual species are represented by grey lines and the average is shown in a pink to yellow gradient. Higher scores are more likely to be a coiled-coil domain with an arbitrary cut off for a coiled-coil shown in a grey dashed line at 0.8. (B) Structural alignment of *Dictyostelium* Skp1A dimer NMR structure (PDB structure 6V88, gray) and *P. pacificus* SKR-1 (teal) with *P. pacificus* residues 20 – 47 and corresponding residues in *Dictyostelium* labeled in blue and black, respectively.

